# Identifying persistent structures in multiscale ‘omics data

**DOI:** 10.1101/2020.06.16.151555

**Authors:** Fan Zheng, She Zhang, Christopher Churas, Dexter Pratt, Ivet Bahar, Trey Ideker

## Abstract

In any ‘omics study, the scale of analysis can dramatically affect the outcome. For instance, when clustering single-cell transcriptomes, is the analysis tuned to discover broad or specific cell types? Likewise, protein communities revealed from protein networks can vary widely in sizes depending on the method. Here we use the concept of “persistent homology”, drawn from mathematical topology, to identify robust structures in data at all scales simultaneously. Application to mouse single-cell transcriptomes significantly expands the catalog of identified cell types, while analysis of SARS-COV-2 protein interactions suggests hijacking of WNT. The method, HiDeF, is available via Python and Cytoscape.

## Background

Significant patterns in data often become apparent only when looking at the right scale. For example, single-cell RNA sequencing data can be clustered coarsely to identify broad categories of cells (e.g. mesoderm, ectoderm), or analyzed more sharply to delineate highly specific subtypes (e.g. pancreas islet β-cells, thymus epithelium) [1–3]. Likewise, protein-protein interaction networks can inform groups of proteins spanning a wide range of spatial dimensions, from protein dimers (e.g. leucine zippers) to larger complexes of dozens or hundreds of subunits (e.g. proteasome, nuclear pore) to entire organelles (e.g. centriole, mitochondria) [4]. Many different approaches have been devised or applied to detect structures in biological data, including standard clustering, network community detection, and low-dimensional data projection [5–7], some of which can be tuned for sensitivity to objects of a certain size or scale (so-called ‘resolution parameters’) [8, 9]. Even tunable algorithms, however, face the dilemma that the particular scale(s) at which the significant biological structures arise are usually unknown in advance.

Guidelines for detecting robust patterns across scales come from the field of topological data analysis, which studies the geometric “shape” of data using tools from algebraic topology and pure mathematics [10]. A fundamental concept in this field is “persistent homology” [11], the idea that the core structures intrinsic to a dataset are those that persist across different scales. Recently, this concept has begun to be applied to analysis of ‘omics data and particularly biological networks [12, 13]. Here, we sought to integrate concepts from persistent homology with existing algorithms for network community detection, resulting in a fast and practical multiscale approach we call the <u>H</u>ierarchical community <u>D</u>ecoding <u>F</u>ramework (HiDeF).

## Results and Discussion

HiDeF works in the three phases to analyze the structure of a biological dataset (**Methods**). To begin, the dataset is formulated as a similarity network, depicting a set of biological entities (e.g. genes, proteins, cells, patients, or species) and pairwise connections among these entities (representing similarities in their data profiles). The goal of the first phase is to detect network communities, i.e. groups of densely connected biological entities. Communities are identified continually as the spatial resolution is scanned, producing a comprehensive pool of candidates across all scales of analysis (**Fig. 1a**). In the second phase, candidate communities arising at different resolutions are pairwise aligned to identify those that have been redundantly identified and are thus *persistent* (**Fig. 1b**). In the third phase, persistent communities are analyzed to identify cases where a community is fully or partially contained within another (typically larger) community, resulting in a hierarchical assembly of nested and overlapping biological structures (**Fig. 1c,d**). HiDeF is implemented as a Python package and can be accessed interactively in the Cytoscape network analysis and visualization environment [14] (**Availability of data and materials**).

**Figure 1.**
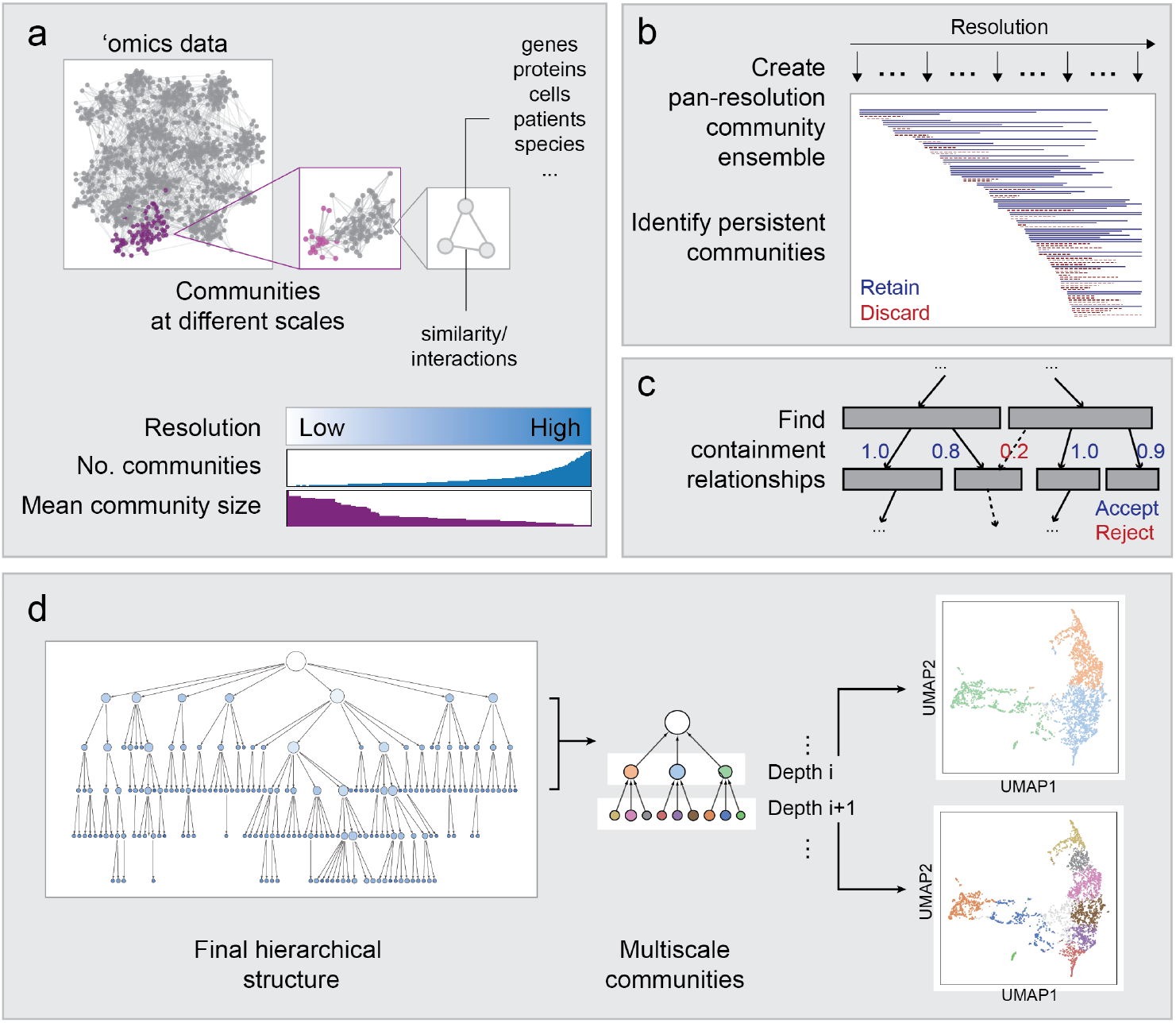
Identification of persistent communities with HiDeF. **a,** ‘Omics data often contain community structures at different spatial resolutions. Increasing the resolution of the analysis generally increases the number of communities and decreases the average community size. **b,** Pan-resolution community detection yields a candidate pool of communities. Communities that are robustly identified across a wide range of resolutions are considered persistent and retained. **c,** Set containment analysis is used to define the relationships between communities, leading to **d,** the final hierarchical model, in which vertices of increasing depths from the root represent communities of increasingly high resolutions.

We first explored the idea of measuring community persistence *via* analysis of synthetic datasets [15] in which communities were simulated and embedded in the similarity network at two different scales (**Supplementary Fig. 1a**; **Methods**). Notably, the communities determined to be most persistent by HiDeF were found to accurately recapitulate the simulated communities at the two scales (**Supplementary Fig. 1b-g**). In contrast, applying community detection algorithms at a fixed resolution had limited capability to capture both scales of simulated structures simultaneously (**Supplementary Fig. 2; Methods**).

We next evaluated whether persistent community detection improves the characterization of cell types. We applied HiDeF to detect robust nested communities within cell-cell similarity networks based on the mRNA expression profiles of 100,605 single cells gathered across the organs and tissues of mice (obtained from two datasets in the *Tabula Muris* project [16]; **Methods**). These cells had been annotated with a controlled vocabulary of cell types from the Cell Ontology (CO) [17], *via* analyses of cell-type-specific expression markers [16]. We used groups of cells sharing the same annotations to define a panel of 136 reference cell types and measured the degree to which each reference cell type could be recapitulated by a HiDeF community of cells (**Methods**). We compared these results to TooManyCells [18] and Conos [19], two recently developed methods that generate nested communities of single cells in divisive and agglomerative manners, respectively (**Methods**). Reference cell types tended to better match communities generated by HiDeF than those of other approaches, with 65% (89/136) having a highly overlapping community (Jaccard index > 0.5) in the HiDeF hierarchy (**Fig. 2a,b, Supplementary Fig. 3a,b**). This favorable performance was observed consistently when adjusting HiDeF parameters to formulate a simple hierarchy, containing only the strongest structures, or a more complex hierarchy including additional communities that are less persistent but still significant (**Fig. 2c, Supplementary Fig. 3c**).

**Figure 2.**
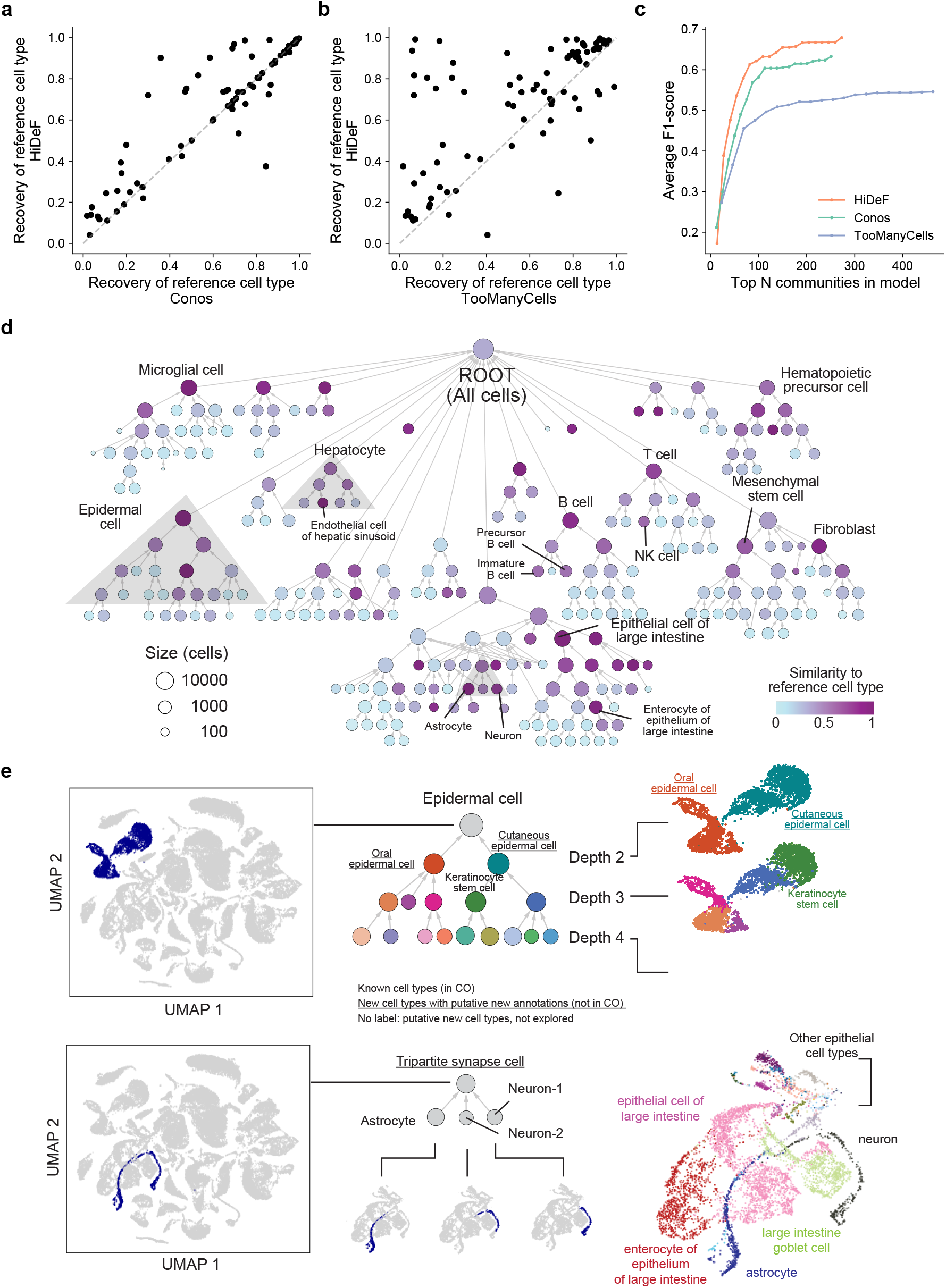
A hierarchy of mammalian cell types from single-cell transcriptomes. **a-b,** Recovery of individual reference cell types by HiDeF (y axis) in comparison to Conos [19] or TooManyCells [18] (x axis of panels a or b, respectively). For each reference cell type (points), the extent of recovery is measured as the maximum F1-score of the set of reference cells with those of any detected community (see **Methods**). **c,** Recovery of reference cell types (evaluated by the average F1-score) among the top *N* ranked cell communities. Communities are ranked in the descending order of score for each community detection tool (e.g. persistence in HiDeF; see **Methods**). **d,** Hierarchy of 273 putative mouse cell types identified by HiDeF. Vertices are cell communities, with color gradient indicating the extent of the optimal match (Jaccard similarity) to a reference cell type. Selected matches to reference cell types are labeled. Gray regions indicate sub-hierarchies (epidermal cells, astrocytes/neurons, and hepatocytes) related to subsequent panels and other figures (**Supplementary Fig. 4**). **e**, Epidermal cell communities. Left: UMAP 2D projection of all cells, with epidermal cells highlighted in dark blue. Middle: Sub-hierarchy of epidermal cell communities as determined by HiDeF. Right: Correspondence between the UMAP projection and the sub-hierarchy, with colors marking the same cell populations across the two representations. **f,** Astrocyte and neuron communities. Left: UMAP 2D projection of all cells, with astrocytes and neurons highlighted in dark blue. Middle: Sub-hierarchy of astrocyte and neuron communities as determined by HiDeF. Cells in the three small communities are highlighted in the below UMAP projections. Right: Broader UMAP context with cells colored and labeled as per the original *Tabula Muris* analysis [16]. Results in this figure are based on the FACS dataset in the *Tabula Muris* [16]; results for the *Tabula Muris* droplet dataset are in **Supplementary Fig. 3**.

The top-level communities in the HiDeF hierarchy corresponded to broad cell lineages such as “T cell”, “B cell”, and “epidermal cell”. Finer-grained communities mapped to more specific known subtypes (**Fig. 2d**) or, more frequently, putative new subtypes within a lineage. For example, “epidermal cell” was split into two distinct epidermal tissue locations, skin and tongue; further splits suggested the presence of still more specific uncharacterized cell types (**Fig. 2e**). HiDeF communities also captured known cell types that were not apparent from 2D visual embeddings (**Supplementary Fig. 4a,b**), and also suggested new cell-type combinations. For example, astrocytes were joined with two communities of neuronal cells to create a distinct cell type not observed in the hierarchies of TooManyCells [18], Conos [19], or a two-dimensional data projection with UMAP [20] (**Fig. 2f, Supplementary Fig. 4c**). This community may correspond to the grouping of a presynaptic neuron, postsynaptic neuron, and a surrounding astrocyte within a so-called “tripartite synapse” [21].

Next, we applied HiDeF to analyze protein-protein interaction networks, with the goal of characterizing protein complexes and higher-order protein assemblies spanning spatial scales. We benchmarked this task by the agreement between HiDeF communities and the Gene Ontology (GO) [22], a database that manually assigns proteins to cellular components, processes, or functions based on curation of literature (**Methods**). Application to protein-protein interaction networks from budding yeast and human found that HiDeF captured knowledge in GO more significantly than previous pipelines proposed for this task, including the NeXO approach to hierarchical community detection [23] and standard hierarchical clustering of pairwise protein distances calculated by three recent network embedding approaches [24–26] (**Fig. 3a,b**; **Supplementary Figs. 5–6**; **Methods**). HiDeF could be directly applied to the original interaction networks or to network embedded versions to further improve the performance and robustness (**Supplementary Fig. 7**).

**Figure 3.**
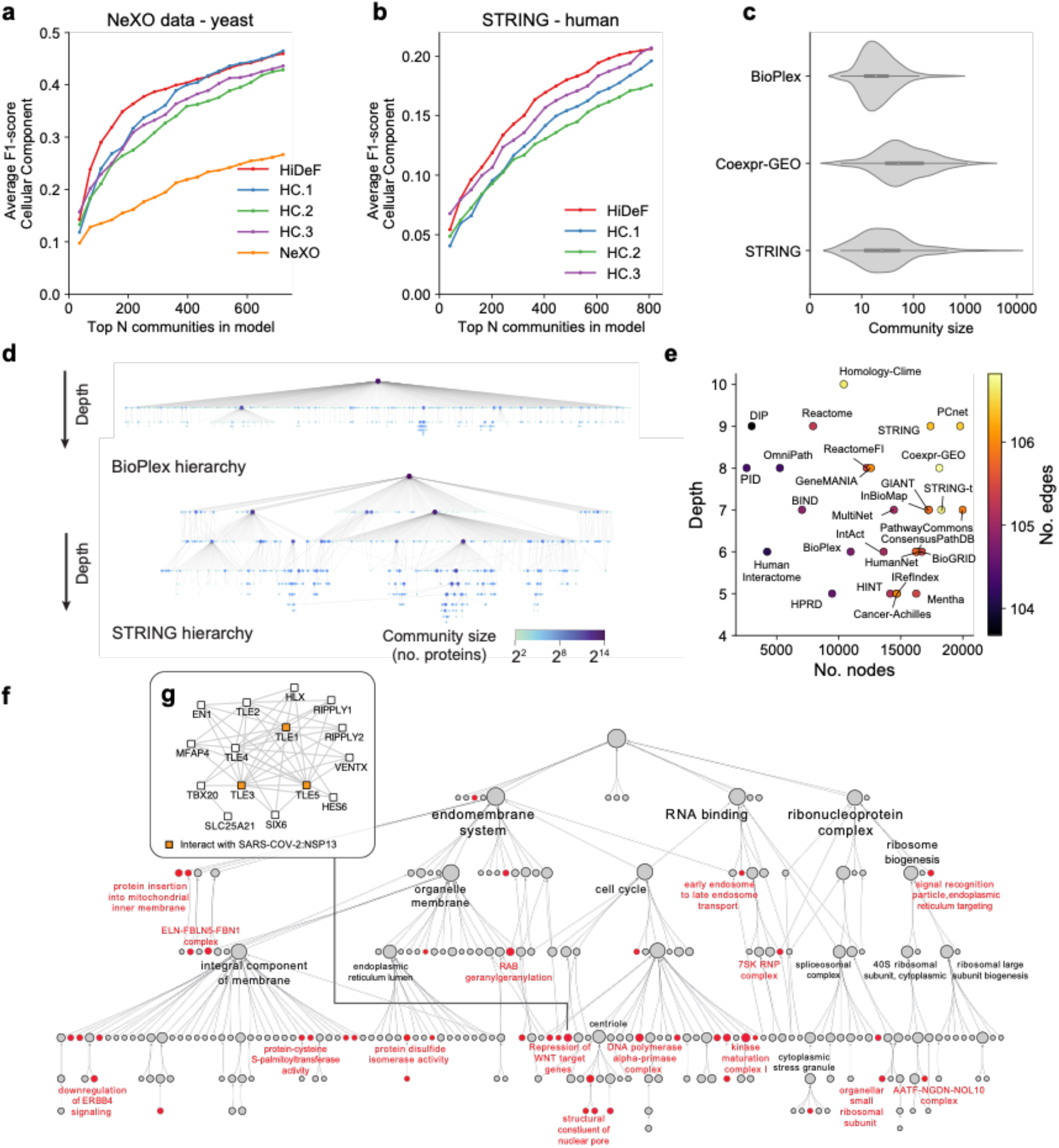
Hierarchical community structure of protein networks. **a-b**, Recovery of cellular components documented in GO by community detection methods (colored traces) versus number of top communities examined. Recovery is evaluated by the average F1 score. Communities are ranked in descending order of score for each community detection tool, similar to **Fig. 2c** (see **Methods**). A yeast network [23] and the human STRING network [31] were used as the inputs of **a** and **b**, respectively. HC.1-3 represent UPGMA <u>H</u>ierarchical <u>C</u>lustering of pairwise distances generated by Mashup, DSD, and deepNF [24–26], respectively. **c,** Distributions of community sizes (x-axis, number of proteins) for three human protein networks: BioPlex 2.0 [29], Coexpr-GEO [30], and STRING [31]. **d,** Community hierarchies identified for BioPlex 2.0 (upper) or STRING (lower) databases. Vertex sizes and colors indicate the number of proteins in the corresponding communities. **e,** Twenty-seven public databases of protein-protein interaction networks were analyzed by HiDeF and profiled by the maximum depths of their resulting hierarchies (y axis), which do not correlate with their total sizes (numbers of proteins, x axis; numbers of edges, color bar). **f**, A hierarchy of communities of human proteins interacting with SARS-COV-2. The hierarchy, generated by HiDeF (**Methods**), contains 252 communities of 1948 human proteins. Communities colored red are enriched (odds ratio > 1.5) for the 332 human proteins interacting with viral proteins of SARS-COV-2. Selected communities are labeled by gene set enrichment function provided in CDAPS (**Availability of data and materials**). **g,** A community of interacting human proteins targeted by the SARS-COV-2 viral protein Nsp13 (**Methods**). Direct interactors of Nsp13 (TLE1, TLE3, TLE5) are shown in orange.

We also applied HiDeF to analyze a collection of 27 human protein interaction networks [27, 28]. We found significant differences in the distributions of community sizes across these networks, loosely correlating with the different measurement approaches used to generate each network. For example, BioPlex 2.0, a network characterizing biophysical protein-protein interactions by affinity-purification mass-spectrometry (AP-MS) [29], was dominated by small communities of 10-50 proteins, whereas a network based on mRNA coexpression [30] tended towards larger-scale communities of >50 proteins. In the middle of this spectrum, the STRING network, which integrated biophysical protein interactions and gene co-expression with a variety of other features [31], contained both small and large communities (**Fig. 3c**). In agreement with the observation above, the hierarchy of BioPlex had a relatively shallow shape in comparison to that of STRING (and other integrated networks including GIANT and PCNet [27, 32]), in which communities across many scales formed a deep hierarchy (**Fig. 3d,e; Availability of data and materials**).

In contrast to clustering frameworks, HiDeF recognizes when a community is contained by multiple parent communities, which in the context of protein-protein networks suggests that the community participates in diverse pleiotropic biological functions. For example, a community corresponding to the MAPK (ERK) pathway participated in multiple larger communities, including RAS and RSK pathways, sodium channels, and actin capping, consistent with the central roles of MAPK signaling in these distinct biological processes [33] (**Supplementary Fig. 8**). The hierarchies of protein communities identified from each of these networks have been made available as a resource in the NDEx database [34] (**Availability of data and materials**).

To explore multiscale data analysis in the context of an urgent public health issue, we considered a recent application of AP-MS that characterized interactions between the 27 SARS-COV-2 viral subunits and 332 human host proteins [35]. We used network propagation to select a subnetwork of the BioPlex 3.0 human protein interactome [36] proximal to these 332 proteins (1948 proteins and 22,835 interactions) and applied HiDeF to identify its community structure (**Methods**). Among the 251 persistent communities identified (**Fig. 3f**), we noted one consisting of human Transducin-Like Enhancer (TLE) family proteins, TLE1, TLE3, and TLE5, which interacted with SARS-COV2 Nsp13, a highly conserved RNA synthesis protein in corona and other nidoviruses (**Fig. 3g**) [37]. TLE proteins are well-known inhibitors of the Wnt signaling pathway [38]. Inhibition of WNT, in turn, has been shown to reduce coronavirus replication [39] and recently proposed as a COVID-19 treatment [40]. If interactions between Nsp13 and TLE proteins can be shown to facilitate activation of WNT, TLEs may be of potential interest as drug targets.

## Conclusions

Community persistence provides a basic metric for distilling biological structure from data, which can be tuned to select only the strongest structures or to include weaker patterns that are less persistent but still significant. This concept applies to diverse biological subfields, as demonstrated here for single cell transcriptomics and protein interaction mapping. While these subfields currently employ very different analysis tools which largely evolve separately, it is perhaps high time to seek out core concepts and broader fundamentals around which to unify some of the ongoing development efforts. To that effect, the methods explored here have wide applicability to analyze the multiscale organization of many other biological systems, including those related to chromosome organization, the microbiome and the brain.

## Methods

### Overview of the approach

Consider an undirected network graph *G*, representing a set of biological *objects* (vertices) and a set of *similarity relations* between these objects (edges). Examples of interest include networks of cells, where edges represent pairwise cell-cell similarity in transcriptional profiles characterized by single-cell RNA-seq, or networks of proteins, where edges represent pairwise protein-protein biophysical interactions. We seek to group these objects into *communities* (subsets of objects) that appear at different scales and identify approximate *containment relationships* among these communities, so as to obtain a hierarchical representation of the network structure. The workflow is implemented in three phases. Phase I identifies communities in *G* at each of a series of spatial resolutions *γ*. Phase II identifies which of these communities are *persistent* by way of a *pan-resolution community graph G_c_*, in which vertices represent communities, including those identified at each resolution, and each edge links pairs of similar communities arising at different resolutions. Persistent communities correspond to large components in *G_c_*. Phase III constructs a final hierarchical structure *H* that represents containment and partial containment relationships (directed edges) among the persistent communities (vertices).

### Phase I: Pan-resolution community detection

Community detection methods generally seek to maximize a quantity known as the *network modularity*, as a function of community assignment of all objects [41]. A *resolution parameter* integrated into the modularity function can be used to tune the scale of the communities identified [9, 42, 43], with larger/smaller scale communities having more/fewer vertices on average (**Fig. 1a**). Of the several types of resolution parameter that have been proposed, we adopted that of the Reichardt-Bornholdt configuration model [42], which defines the generalized modularity as:

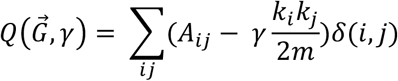

where 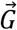 defines a mapping from objects in *G* to community labels; *k_i_* is the degree of vertex *i*; *m* is the total number of edges in *G*; *γ* is the resolution parameter; *δ*(*i,j*) indicates that vertices *i*. and *j* are assigned to the same community by 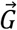; and *A* is the adjacency matrix of *G*. To determine 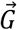 we use the extended Louvain algorithm implemented in the Python package louvain-igraph (http://github.com/vtraag/louvain-igraph; version 0.6.1). Values of *γ* are sampled logarithmically between lower and upper bounds *γ_min_* and *γ_max_* at a minimum density such that for all *γ* there exist at least 10 nearby *γ′* satisfying:

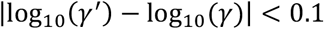

Two *γ* values satisfying the above formula are defined as *γ-proximal*. The sampling step, which was practically set to 0.1 to sufficiently capture the interesting structures in the data; it is conceptually similar to the Nyquist sampling frequency in signal processing [44]. We used *γ_min_* = 0.001, which we found always resulted in the theoretical minimum number of communities, equal to the number of connected components in *G*. We used *γ_max_* = 20 for single-cell data (**Fig. 2, Supplementary Fig. 3, 4**) and *γ_max_* = 50 for simulated networks (**Supplementary Figs. 1, 2**) and protein interaction networks (**Fig. 3, Supplementary Figs. 5–8**). Performing Louvain community detection at each *γ* over this defined progression of values resulted in a set of communities 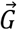 at each *γ*.

### Phase II: Identification of persistent communities

To identify persistent communities, we define the pairwise similarity between any two communities *a* and *b* as the Jaccard similarity of their sets of objects, *s*(*a*) and *s*(*b*):

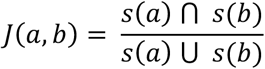

Pairwise community similarity is computed only for pairs of communities identified at two different *γ*-proximal resolution values, as communities within a resolution do not overlap. To represent these similarities, we define a *pan-resolution community graph G_c_*, in which vertices are communities identified at any resolution and edges connect pairs of similar communities having *J*(*a, b*) > *τ*. Each component of *G_c_* defines a family of similar communities spanning resolutions, for which the *persistence* can be naturally defined by the number of distinct *γ* values covered by the component. For each component in *G_c_* larger than a persistence threshold *χ*, the biological objects participating in more than *p* % of communities represented by the vertices of that component define a *persistent community*.

### Phase III: A hierarchy of nested and overlapping communities

We initialize a hierarchical structure represented by *H*, a directed acyclic graph (DAG) in which each vertex represents a persistent community. A *root vertex* is added to represent the community of all objects. The containment relationship between two vertices, *v* and *w*, is quantified by the *containment index* (*CI*):

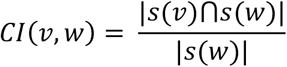

which measures the fraction of objects in *w* shared with *v*. An edge is added from *v* to *w* in *H* if *CI*(*v,w*) is larger than a threshold *σ* (*w* is *σ-contained* by *v*). Since *J*(*v,w*) < *τ* for all *v, w* (a property established by the procedure for connecting similar communities in phase II), setting *σ* ≥ 2*τ*/(1 + *τ*) guarantees *H* to be acyclic. In practice we used a relaxed threshold *σ* = *τ*, which we found generally maintains the acyclic property but includes additional containment relations. In the (in our experience rare) event that cycles are generated in *H*, i.e. *CI*(*v,w*) ≥ *τ* and *CI*(*w, v*) ≥ *τ*, we add a new community to *H*, the union of *v* and *w*, and remove *v* and *w* from *H*.

Finally, redundant relations are removed by obtaining a transitive reduction [45] of *H*, which represents the hierarchy returned by HiDeF describing the organization of communities. The biological objects assigned to each community are expanded to include all objects assigned to its descendants. Throughout this study, we used the parameters *τ* = 0.75, *χ* = 5, *p* = 75. Note that since *χ* is a threshold of minimum persistence, the results under a larger value of *χ′* can be produced by simply removing communities with persistence lower than *χ′* (**Figs. 2c, 3a-b; Supplementary Figs 2, 3c, 5**). Generally, we observed that the conclusions drawn in this study were robust to this choice of parameters. The persistence of communities only moderately correlates with community sizes, with the consequence that different choices of persistent threshold *χ* do not strongly favor structures at particular scales. (**Supplementary Fig. 9**). Different combinations of parameters *τ* and *p* typically do not significantly change the performance of HiDeF in the benchmark tests on protein-protein interaction networks (**Supplementary Fig. 6**), except that certain parameters (e.g. *τ* = 0.9) are less robust to network perturbation (i.e. randomly deleting edges from networks). We found that combining HiDeF with node embedding resolved this issue and further improved the performance and robustness (**Supplementary Fig. 7;** see sections below).

### Simulated benchmark networks

Simulated network data were generated using the Lancichinetti-Fortunato-Radicchi (LFR) method [15] (**Supplementary Figs. 1, 2**). We used an available implementation (LFR benchmark graphs package 5 at http://www.santofortunato.net/resources) to generate benchmark networks with two levels of embedded communities, a coarse-grained (macro) level and a fine-grained (micro) level. Within each level, a vertex was exclusively assigned to one community. Two parameters, *μ*_c_ and *μ*_f_, were used to define the fractions of edges violating the simulated community structures at the two levels. All other edges were restricted to occur between vertices assigned to the same community (**Supplementary Fig. 1a**). We fixed other parameters of the LFR method to values explored by previous studies [9]. In particular, *N* = 1000 (number of vertices), *k* = 10 (average degree), *maxk* = 40 (maximum degree), *minc* = 5 (minimum number of vertices for a micro-community), *maxc* = 20 (maximum number of vertices for a microcommunity), *minC* = 50 (minimum number of vertices for a macro-community), *maxC* = 100 (maximum number of vertices for a macro-community), *t_1_* = 2 (minus exponent for the degree sequence), *t_2_* = 1 (minus exponent for the community size distribution). The numbers of coarsegrained communities and fine-grained communities in each simulated network were approximately bounded by *minC*, *maxC*, *minc* and *maxc* (10-20 and 50-200, respectively), and the sizes of communities within each level were set to be close to each other (as *t_2_* = 1).

Some community detection algorithms include iterations of local optimization and vertex aggregation, a process that, like HiDeF, also defines a hierarchy of communities, albeit as a tree rather than a DAG. We demonstrated that without scanning multiple resolutions, this process alone was insufficient to detect the simulated communities at all scales (**Supplementary Fig. 2**). We used Louvain and Infomap [46, 47], which have stable implementations and have shown strong performance in previous community detection studies [48]. For Louvain, we optimized the standard Newman-Girvan modularity (equivalent to *γ* = 1, see above) using the implementation at http://github.com/vtraag/louvain-igraph. For Infomap, we used the version 1.0.0-beta.47 from https://www.mapequation.org/, and set ‘Markov time’ (the ‘resolution parameter’ of Infomap) to 1 and other parameters to default. In general, these settings generated trees with two levels of communities. Note that Infomap sometimes determined that the input network was non-hierarchical, in which cases the coarse- and fine-grained communities were identical by definition.

### Single-cell RNA-seq data

Mouse single-cell RNA-seq data (**Fig. 2; Supplementary Fig. 3**) were obtained from the Tabula Muris project [16], (https://tabula-muris.ds.czbiohub.org/;), which contains two datasets generated with different experimental methods of separating bulk tissues into individual cells: FACS and microfluidic droplet. We applied HiDeF to the shared nearest neighbor graph parsed from the data files (R objects; accessible at https://doi.org/10.6084/m9.figshare.5821263.v2) provided in that study. All data normalization and pre-processing procedures have been described in the *Tabula Muris* paper [16]. Briefly, counts were log-normalized using the natural logarithm of 1 + counts per million (for FACS) or 1 + counts per ten thousand (for droplet). A threshold (0.5) for the standardized log dispersion was used to select variable genes. A shared nearest neighbor (SNN) graph was then created by the Seurat *‘FindNeighbors’* function [3] using the first 30 principal components of each dataset.

Identical analyses were applied to the FACS and the droplet datasets respectively, yielding a hierarchy of 273 and 279 communities respectively (**Fig. 2d**). ScanPy 1.4.5 [49] was used to create tSNE or UMAP embeddings and associated two-dimensional visualizations [20] as baselines for comparison (**Fig. 2e,f; Supplementary Fig. 3a,b**). Through previous analysis of the single-cell RNA data, all cells in these datasets had been annotated with matching cell-type classes in the Cell Ontology (CO) [17]. Before comparing these annotations with the communities detected by HiDeF, we expanded the set of annotations of each cell according to the CO structure, to ensure the set also included all of the ancestor cell types of the type that was annotated. For example, CO has the relationship “[keratinocyte] (is_a) [epidermal_cell]”, and thus all cells annotated as “keratinocyte” are also annotated as “epidermal cell”. The CO was obtained from http://www.obofoundry.org/ontology/cl.html and processed by the Data Driven Ontology Toolkit (DDOT) [50] retaining “is_a” relationships only.

We compared HiDeF to TooManyCells [18] and Conos [19] as baseline methods. The former is a divisive method which iteratively applies bipartite spectral clustering to the cell population until the modularity of the partition is below a threshold; the latter uses the Walktrap algorithm to agglomeratively construct the cell-type hierarchy [51]. We chose to compare with these methods because their ability to identify multiscale communities was either the main advertised feature or had been shown to be a major strength. TooManyCells (version 0.2.2.0) was run with the parameter *“*min-modularity*”* set to 0.025 as recommended in the original paper [18], with other settings set to default. This process generated dendrograms (binary trees) with 463 communities. The Walktrap algorithm was run from the Conos package (version 1.2.1) with the parameter “step” set to 20 as recommended in the original paper [19], yielding a dendogram. The *greedyModularityCut* method in the Conos package was used to select *N* fusions in the original dendrogram, resulting in a reduced dendrogram with 2*N*+1 communities (including *N* internal and *N+1* leaf nodes). Here we used *N* = 125, generating a hierarchy with 251 communities (**Fig. 2c**).

The communities in each hierarchy were ranked to analyze the relationships between celltype recovery and model complexity (**Fig. 2c, Supplementary Fig. 3c**). HiDeF communities were ranked by their persistence; Conos and TooManyCells communities were ranked according to the modularity scores those methods associate with each branch-point in their dendrograms. Conos/Walktrap uses a score based on the gain of modularity in merging two communities, whereas TooManyCells uses the modularity of each binary partition.

### Protein-protein interaction networks

We obtained a total of 27 human protein interaction networks gathered previously by survey studies [27, 28], along with one integrated network from budding yeast (*S. cerevisiae*) that had been used in a previous community detection pipeline, NeXO [23]. This collection contained two versions of the STRING interaction database, with the second removing edges from text mining (labeled STRING-t versus STRING, respectively; **Fig. 3**). Benchmark experiments for the recovery of the Gene Ontology (GO) were performed with STRING and the yeast network (**Fig. 3a,b**, **Supplementary Fig. 4**). The reference GO for yeast proteins was obtained from http://nexo.ucsd.edu/. A reference GO for human proteins was downloaded from http://geneontology.org/ via an API provided by the DDOT package [50].

HiDeF was directly applied to all of the above benchmark networks. The NeXO communities were obtained from http://nexo.ucsd.edu/, with a robustness score assigned to each community. To benchmark communities created by hierarchical clustering, we first calculated three versions of pairwise protein distances (HC.1-3; **Fig. 3a,b; Supplementary Fig. 4**) using Mashup, DSD and deepNF [24–26]. Mashup was used to embed each protein as a vector, with 500 and 800 dimensions for yeast and human, as recommended in the original paper. A pairwise distance was computed for each pair of proteins as the cosine distance between the two vectors. Similarly, deepNF was used to embed each protein into a 500-dimensional vector by default. DSD generates pairwise distances by default. Given these pairwise distances, UPGMA clustering was applied to generate binary hierarchical trees. Following the procedure given in the NeXO and Mashup papers [23, 24] communities with <4 proteins were discarded.

Since all methods had slight differences in the resulting number of communities, communities from each method were sorted in decreasing order of score, enabling comparison of results across the same numbers of top-ranked communities. HiDeF communities were ranked by persistence. NeXO communities were ranked by the robustness value assigned to each community in the original paper [23]. To rank each community *c* of hierarchical clustering (branch in the dendrogram), a one-way Mann-Whitney U-test was used to test for significant differences between two sets of protein pairwise distances: (set 1) all pairs consisting of a protein in *c* and a protein in the sibling community of *c*; (set 2) all pairs consisting of a protein in each of the two children communities of *c*. The communities were sorted by the one-sided p-value of significance that distances in set 1 are greater than those in set 2.

### Metric for evaluating the performance of multiscale structure identification

We adopted a metric average F1-score [52] to evaluate the overall performance of multiscale structure identification, focusing on the recovery of reference communities. Given a set of reference communities *C*^*^ and a set of computationally detected communities 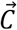, the score was defined as:

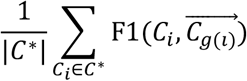

where *g*(*i*) is the best match of *C_i_* in 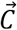, defined as follows:

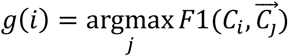

and 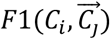 is the harmonic mean of Precision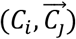 and Recall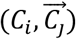. The calculations were conducted by the *xmeasures* package (https://github.com/eXascaleInfolab/xmeasures) [53].

### Combining HiDeF with network embedding

HiDeF was directly applied to the original networks in in most of our analyses of protein-protein interaction networks, and compared with the results of hierarchical clustering following the network embedding techniques [24, 26]. We sought to explore if we can combine the strength of network embedding and HiDeF to further improve the performance and robustness to parameter choices (**Supplementary Fig. 7**). We borrowed the idea of shared-nearest neighbor (SNN) graph that we had been using in the analyses of single-cell data. We made a customized script to use the 500-dimensional node embeddings of the STRING network as the input of the Seurat *FindNeighbors* function [3]. The parameters of this function remained as the default. The output SNN graph has 1.65 × 10^6^ edges, which is on the same magnitude as the original network (2.23 × 10^6^ edges). We then applied HiDeF to this SNN graph with different combinations of parameters (**Supplementary Fig. 7**).

### Analysis of SARS-COV-2 viral-human protein network

332 human proteins identified to interact with SARS-COV-2 viral protein subunits were obtained from a recent study [35]. This list was expanded to include additional human proteins connected to two or more of the 332 virus-interacting human proteins in the new BioPlex 3.0 network [36]. These operations resulted in a network of 1948 proteins and 22,835 interactions. HiDeF was applied to this network with the same parameter settings as for other protein-protein interaction networks (see previous Methods sections), and enrichment analysis was performed via g:Profiler [54] (Fig. 3f,g).

## Declarations

### Ethics approval and consent to participate

Not applicable.

### Consent for publication

Not applicable.

### Availability of data and materials

HiDeF is available through CDAPS (Community Detection APplication and Service), which enables simultaneous visualization of the hierarchical model and the underlying network data and is integrated with the Cytoscape visualization and analysis environment. The Cytoscape App can be downloaded at: http://apps.cytoscape.org/apps/cycommunitydetection. HiDeF is separately available as a Python package: https://github.com/fanzheng10/HiDeF. The version of source code used in this manuscript is deposited in Zenodo: https://doi.org/10.5281/zenodo.4059074.

The hierarchical models generated in this study can be obtained as a network collection within the Network Data Exchange (NDEx) database [34]: https://doi.org/10.18119/N9ZP58. These models include the hierarchy of murine cell types (**Fig. 2**), the hierarchies of yeast and human protein communities identified through protein network analysis, and the hierarchy of human protein complexes targeted by SARS-COV2 (**Fig. 3**).

### Author Contributions

F.Z. designed the study and developed the conceptual ideas. F.Z. and S.Z. implemented the main algorithm. F.Z. and D.P collected the input data and conducted analysis. S.Z. made the Python package. C.C. developed the server that provides the Cytoscape integration. F.Z. and T.I. wrote the manuscript with suggestions from S.Z., D.P. and I.B.

## Acknowledgments

We are grateful for the helpful discussions with Drs. Jianzhu Ma, Karen Mei, and Daniel Carlin.

## Supplementary Figures

**Supplementary Figure 1.**
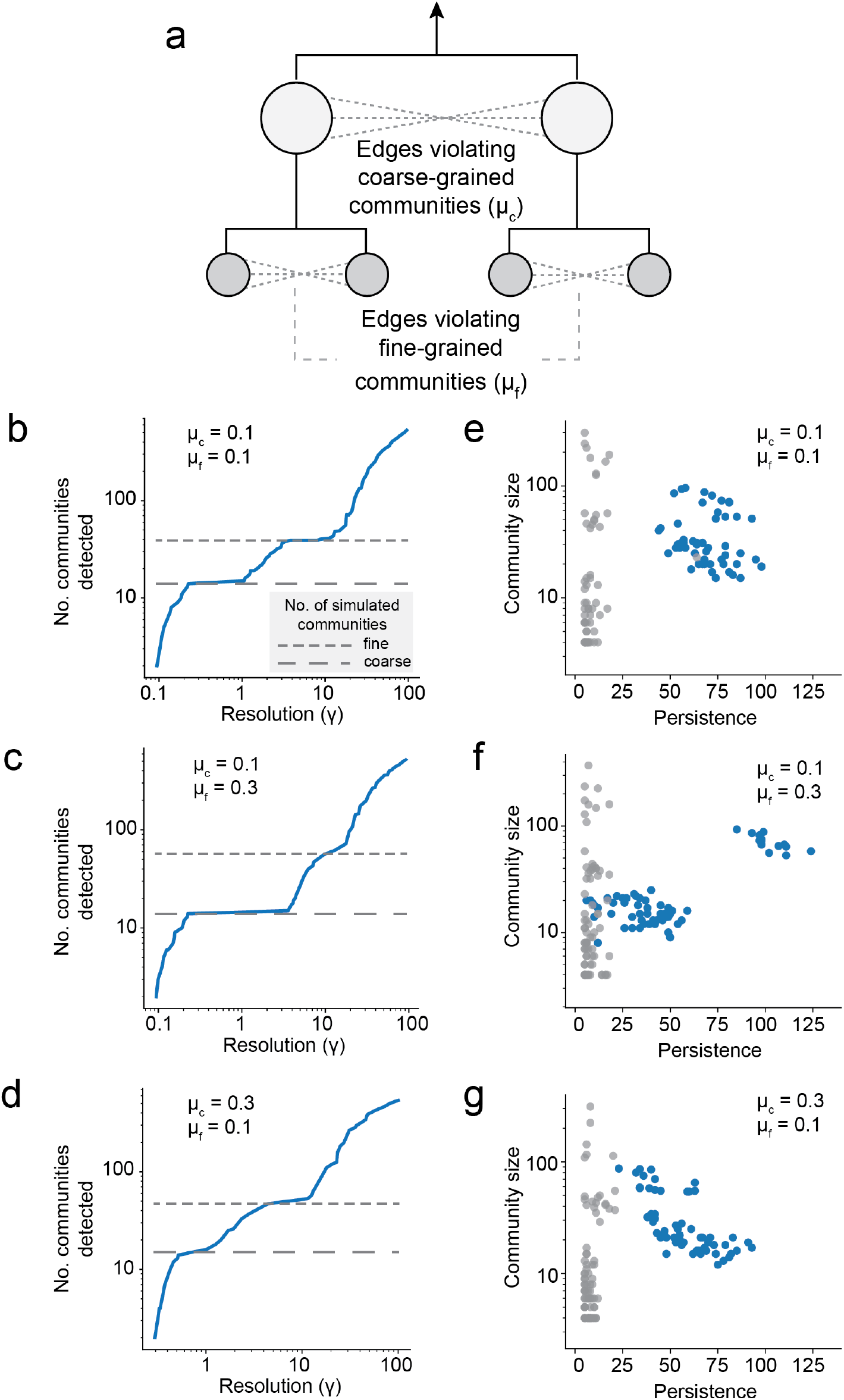
Exploring simulated networks. **a,** The LFR generative model [15] was used to simulate networks with 1000 vertices and average degree 10 (**Methods**). The simulation included two layers of communities, “coarse” (10-20 communities, 50-100 vertices per community) and “fine” (25-200 communities, 5-40 vertices per community), with each fine community nested within a coarse community. Two “mixing parameters” *μ*_c_ and *μ*_F_ controlled the amount of noise, by setting the fraction of edges violating the coarse and fine community structures, respectively. **b-d,** HiDeF analysis of three simulated networks created with different mixing parameters: low balanced noise (b); increased noise in fine communities (c); and increased noise in coarse communities (d). Each plot shows the number of identified communities (y axis) as the resolution is progressively scanned (x axis). The number of communities increases with the resolution parameter, with plateaus matching the actual numbers of coarse and fine communities in the simulated network (dashed lines). Note that the sizes of the plateaus (i.e. the extent of community “persistence”, see text) are affected by the mixing parameters. **e-g**, Companion plots to panels (b-d). Points represent identified communities, delineated by size (y axis) and persistence (x axis). Blue/gray point colors indicate a match/non-match to a true community in the simulated network (Jaccard similarity > 0.75). Note that when noise is low (e), the highest persistence communities correctly recover simulated communities with near-perfect accuracy, e.g. for persistence threshold >20.

**Supplementary Figure 2.**
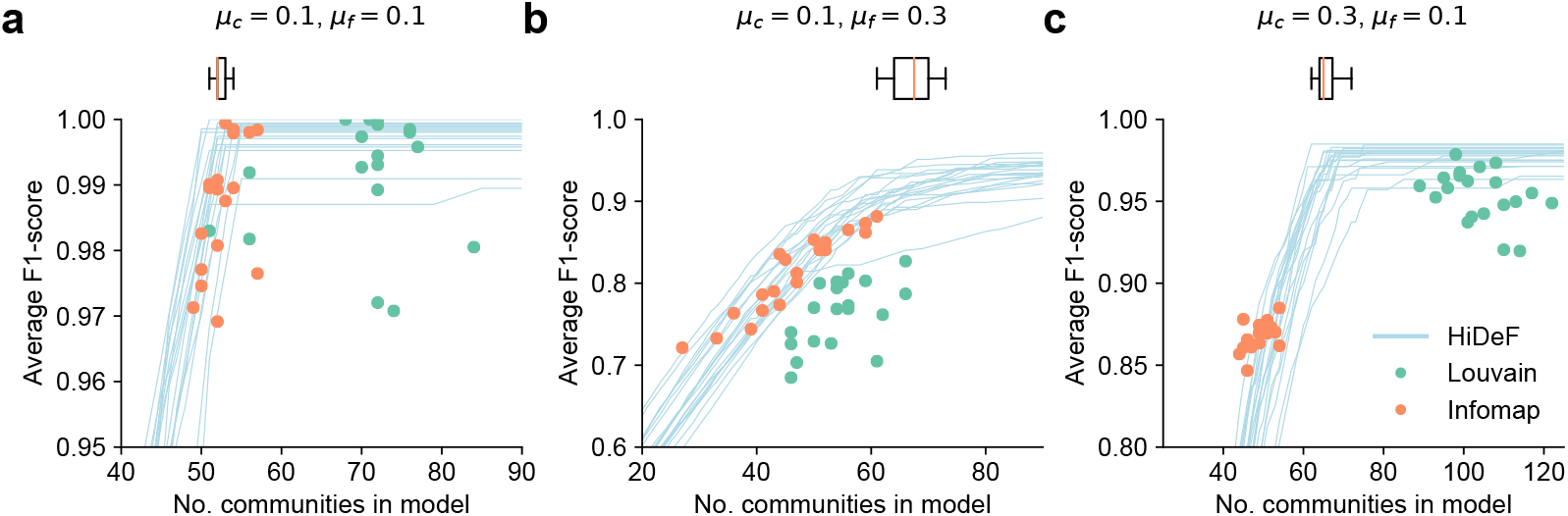
Comparison of methods in recovery of simulated communities. HiDeF is compared with the Louvain and Infomap algorithms [46, 47], with Louvain and Infomap fixed at their default single resolutions (**Methods**). The three plots (**a-c**) compare the performance of the three algorithms in recovering simulated communities at different settings of the coarse/fine mixing parameters (see **Supplementary Fig. 1**). The communities returned by HiDeF are ordered by persistence to evaluate the recovery of among the top *N* most persistent communities (by the average F1-score), whereas Louvain and Infomap generate results with fixed number of communities (green and orange points, respectively). The box plots on the top indicate the numbers of simulated communities. Each plot represents the results for 20 simulations. Note that HiDeF reached the maximum recovery when considering communities at a threshold equal to the number of simulated communities. Louvain and Infomap usually did not generate correct number of communities, and/or generate communities with worse agreements to simulated communities.

**Supplementary Figure 3.**
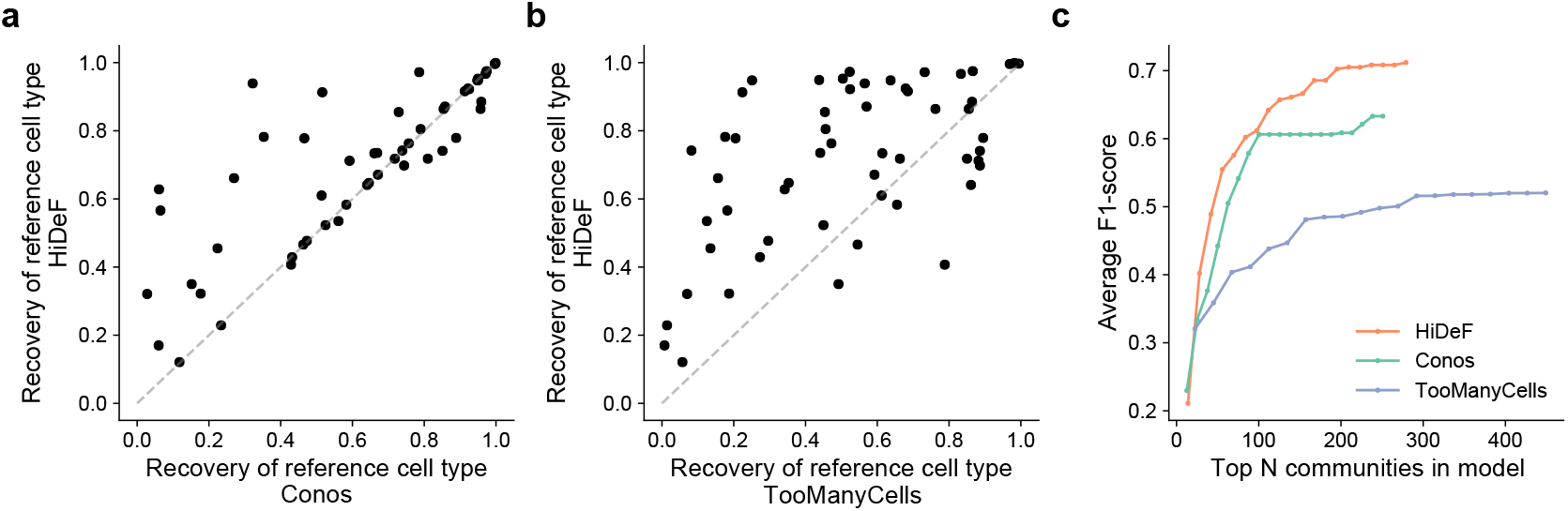
Recovery of mammalian reference cell types from single-cell transcriptomes using the *Tabula Muris* droplet dataset [16]. Similar to **Fig. 2a-c**. **a-b,** Recovery of individual reference cell types by HiDeF (y axis) in comparison to Conos [19] or TooManyCells [18] (x axis of panels a or b, respectively). For each reference cell type (points), the extent of recovery is measured as the maximum F1-score of the set of reference cells with those of any detected community (see **Methods**). **c,** Reference cell types recovered (evaluated by the average F1-score) among the top *N* ranked cell communities. Communities are ranked in the descending order of score for each community detection tool (**Methods**).

**Supplementary Figure 4.**
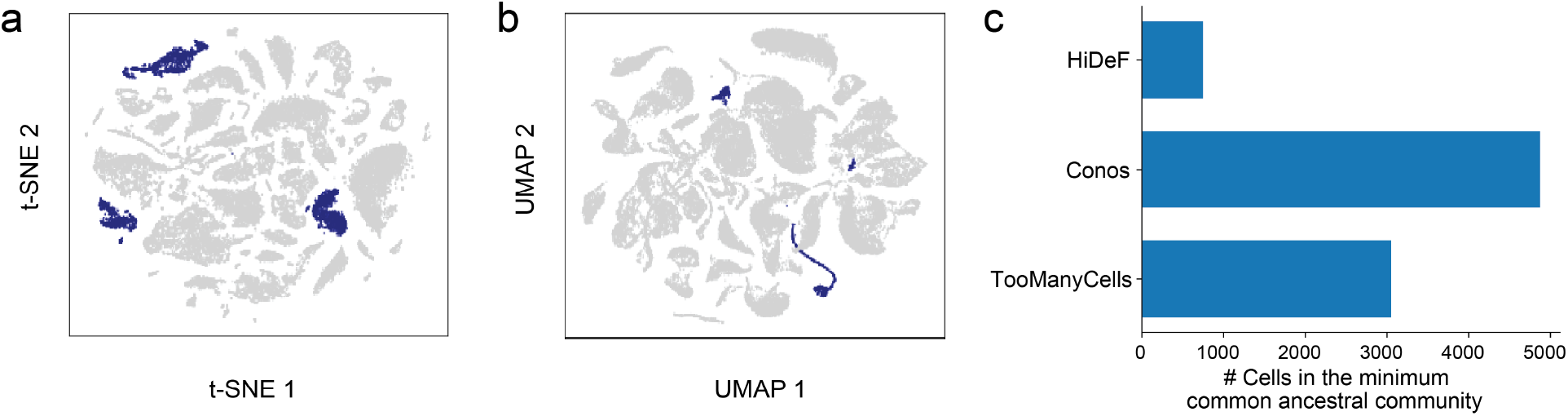
Example cell types captured by HiDeF but not by other approaches. **a,** t-SNE projection of all cells, with the epidermal cell type highlighted (blue). **b,** UMAP projection of all cells, with the hepatocyte cell type highlighted (blue). **c**. Distances between astrocyte and neuron communities in the cell-type hierarchies generated by HiDeF, Conos, or TooManyCells. HiDeF identifies a specific super-community joining both cell types (<1000 cells), whereas such a specific community is not identified by Conos and TooManyCells.

**Supplementary Figure 5.**
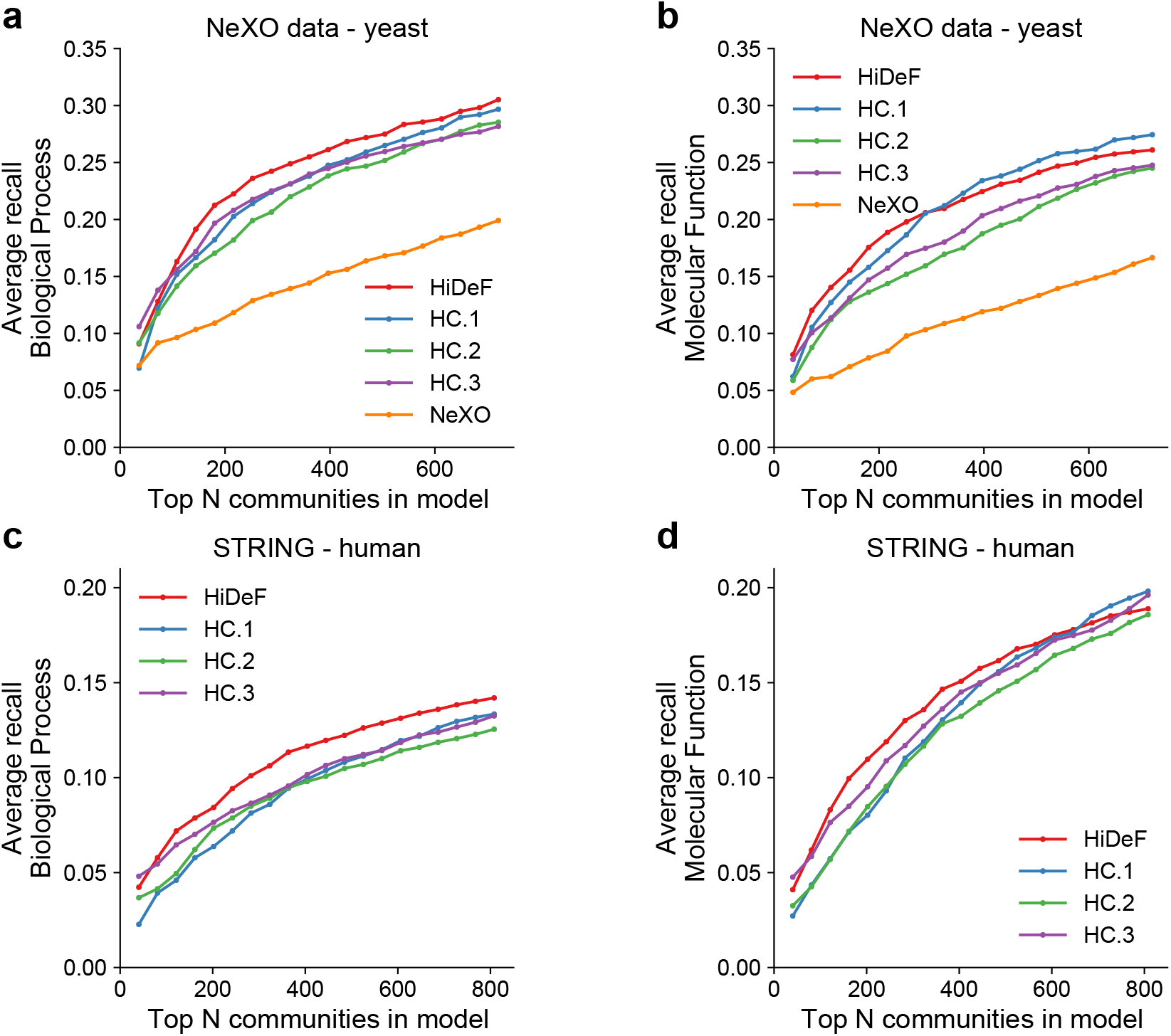
Recovery of GO terms from community detection in protein networks. Similar to **Fig. 3a-b**. HiDeF and alternative methods were applied to build a hierarchy of protein communities from analysis of an integrated protein interaction network for budding yeast (Top: NeXO) or human (Bottom: STRING). The hierarchy of each method (colors) is scored by its recovery of GO terms (average F1 score; Left: Biological Process; Right: Molecular Function) as a function of the number of top-scoring protein communities examined. HC, Hierarchical Clustering following any of three protein pairwise distance functions (Mashup, DSD, and deepNF) [24–26].

**Supplementary Figure 6.**
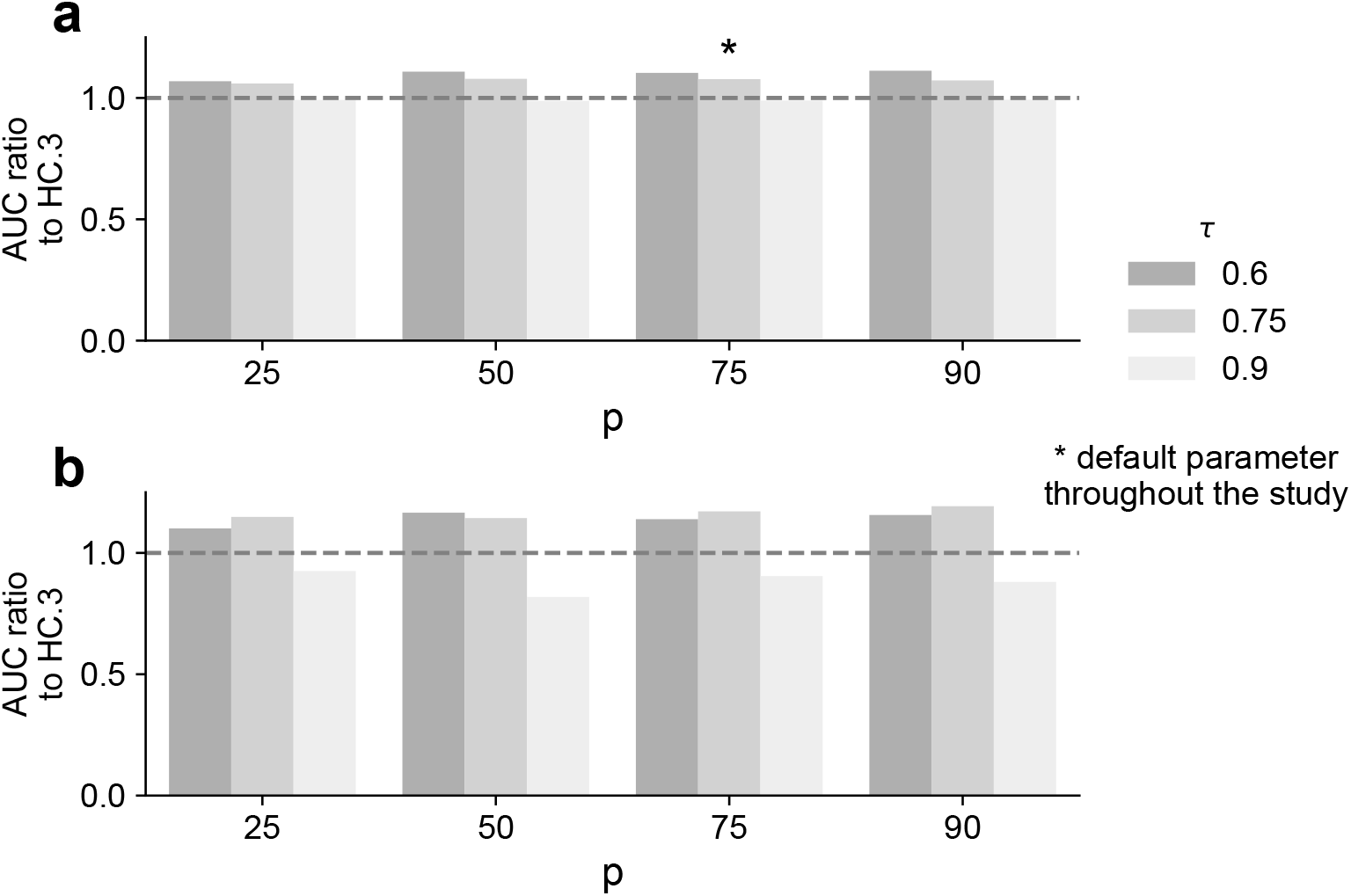
Robustness of GO term recovery to the choice of parameters. **a**, Using the performance analysis depicted in **Fig. 3b**, the Area Under Curve (AUC) was computed for different sets of HiDeF parameters (*p, τ*). This AUC was compared to that of the best baseline tool, HC.3 (i.e. hierarchical clustering of pairwise distances generated by deepNF [26]) to generate an equal number of communities (**Methods**). Note the ratio HiDeF AUC / HC.3 AUC is usually higher than 1, indicating the favorable performance of HiDeF except for very high values of the τ parameter. As per **Fig. 3b**, the analysis was undertaken using the STRING network and the GO Cellular Component branch. **b**, Similar analysis with subsampling of network edges (in which a random 10% of network edges are removed prior to community detection at each resolution).

**Supplementary Figure 7.**
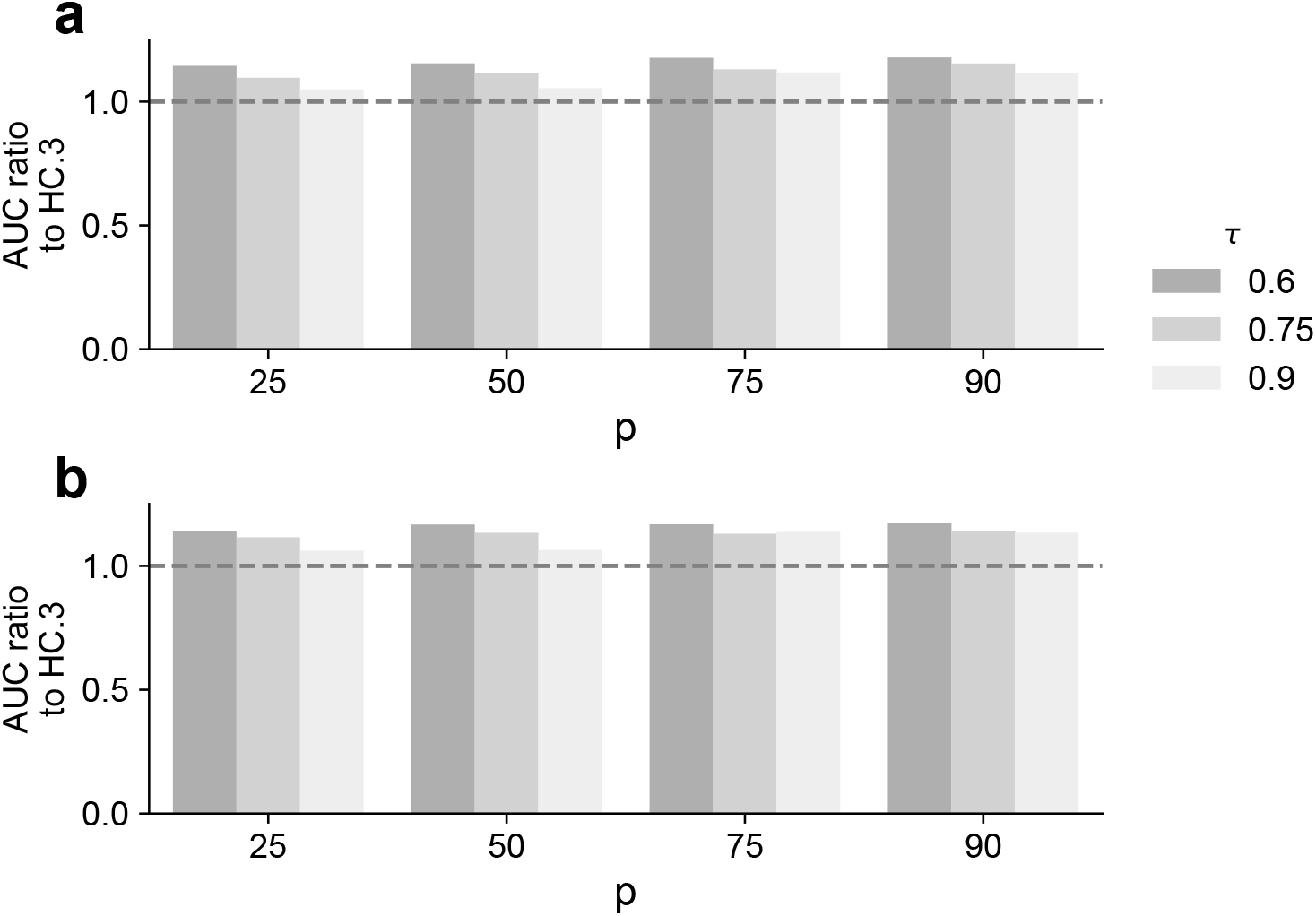
Combining HiDeF and network embedding further improves recovery of GO terms. HiDeF was applied to an SNN graph based on the deepNF embedding (see **Methods**). Other settings of this analysis are identical to that in Supplementary Fig. 6. Note that the performance of recovering GO terms in the Cellular Component branch is now better than HC.3 under all tested parameter settings of HiDeF.

**Supplementary Figure 8.**
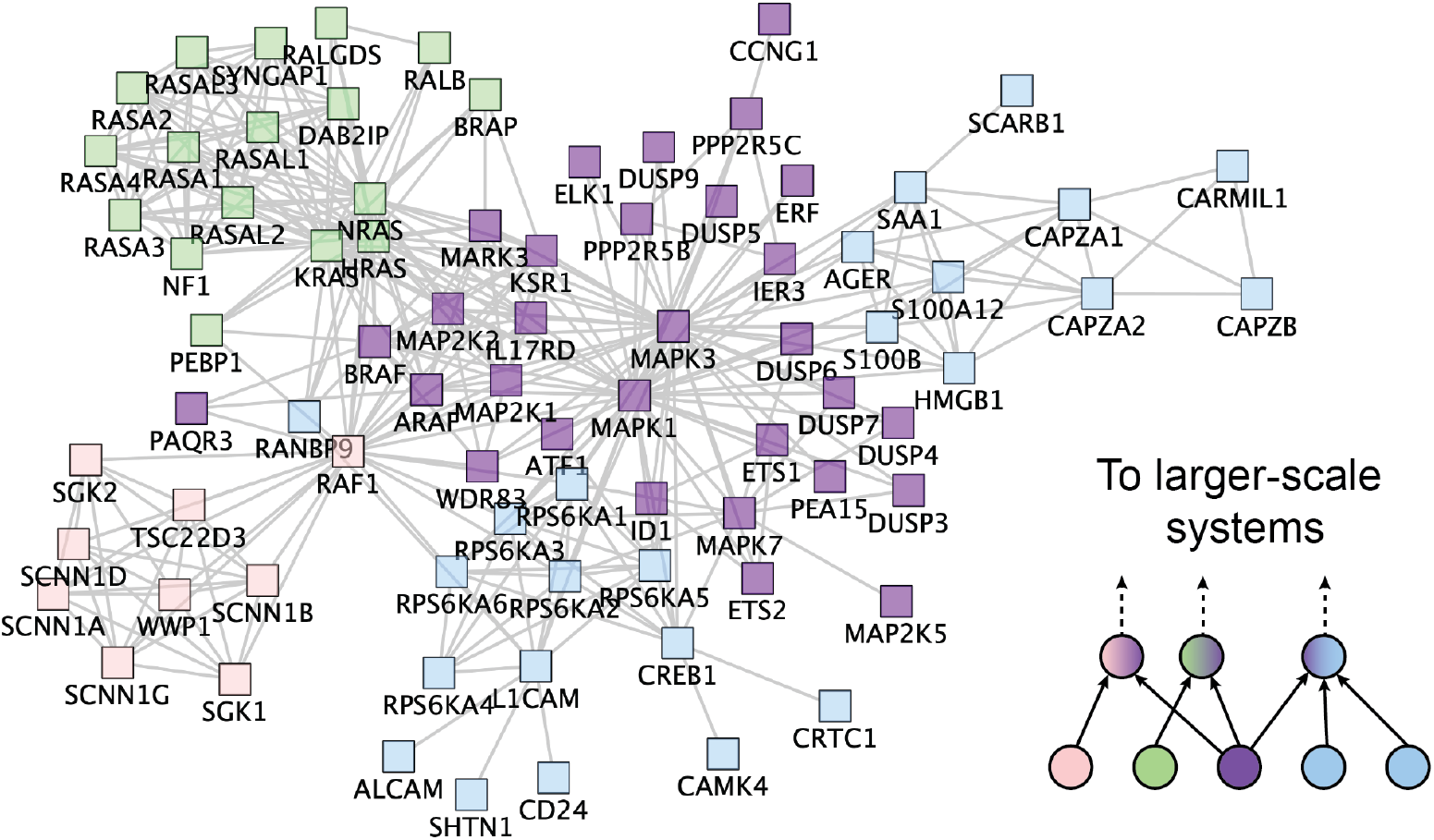
Convergence of communities into multiple super-systems. A community of mitogen-activated protein kinases and dual-specificity phosphatases (purple, center) participates in three distinct larger communities involving separate functions related to RAS pathways (green), sodium channels (pink), and acting capping (blue). The corresponding hierarchical relationships of these communities are depicted at lower right. The source network is Reactome [55].

**Supplementary Figure 9.**
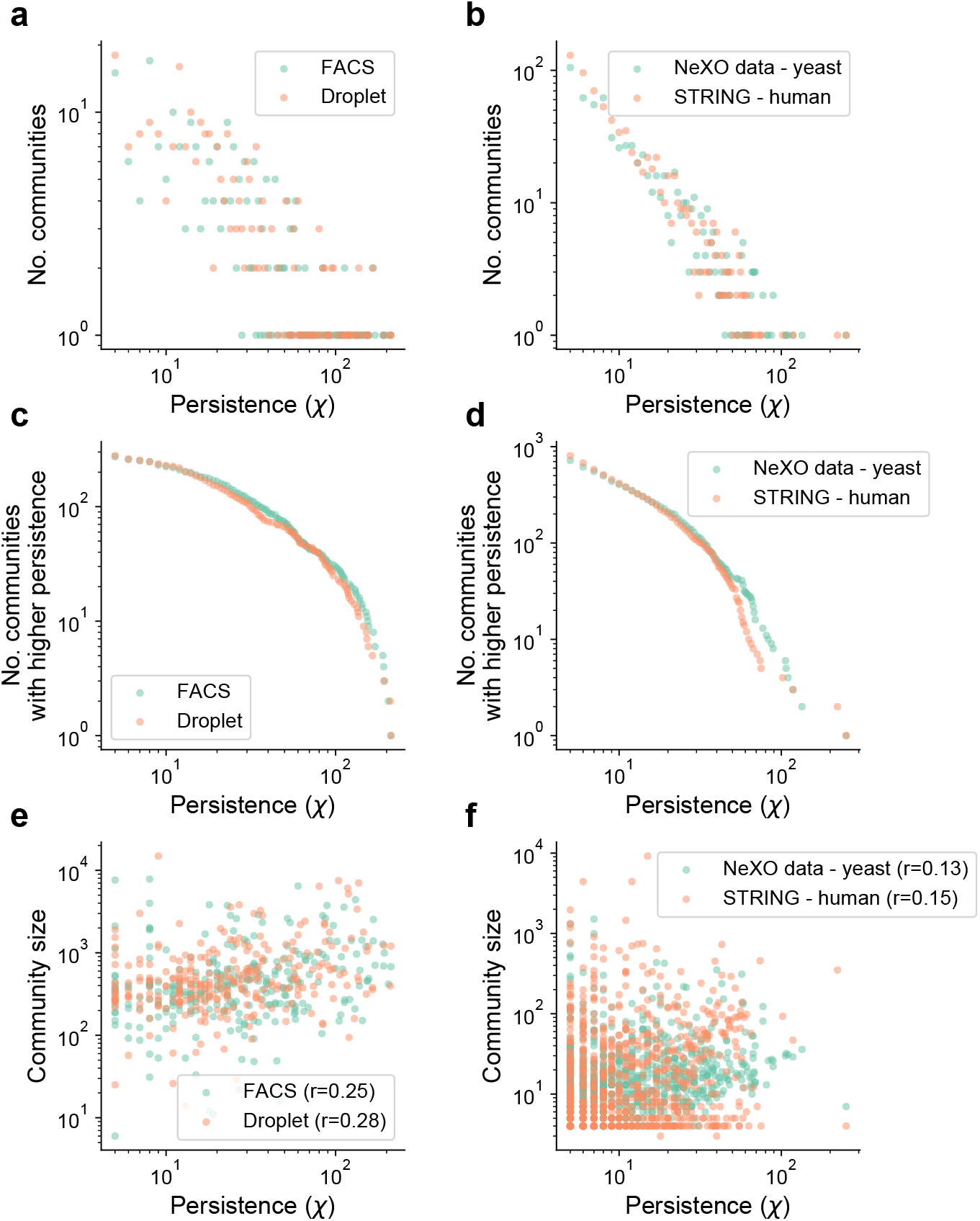
Persistence of HiDeF communities. **a-b,** The number of communities (y axis) at each value of persistence (x axis). **c-d,** The number of communities with higher persistence (y axis) than a given threshold (x axis). **e-f,** Scatterplots of community size (y axis) versus persistence (x axis). The left column characterizes the single-cell transcriptomics data (**Fig. 2, Supplementary Fig. 3**). The right column (panel b, d, f) characterizes the yeast and human protein-protein interaction datasets (**Fig. 3a-b**).

